# Repetitive Somatosensory Stimulation Shrinks The Body Image

**DOI:** 10.1101/2024.06.24.600394

**Authors:** Malika Azaroual-Sentucq, Silvia Macchione, Luke E. Miller, Eric Koun, Romeo Salemme, Matthew R. Longo, Dollyane Muret, Alessandro Farnè

## Abstract

Current models of mental body representations (MBRs) indicate that tactile inputs feed several of them for different functions, implying that altering tactile inputs may affect MBRs differently. Here we tested this hypothesis by leveraging Repetitive Somatosensory Stimulation (RSS), known to improve tactile perception by modulating primary somatosensory cortex (SI) activity, and measured its effects over the *body image*, the *body model* and the *superficial schema* in a randomized sham-controlled, double-blind cross-over study. Results show that RSS affected the *body image*, participants perceiving their finger size as being smaller after RSS. While previous work showed increase of finger size perception after tactile anesthesia (Gandevia & Phegan 1999), these findings reveal that tactile inputs can diametrically modulate the body image. In contrast, RSS did not alter the *body model* or *superficial schema*. In addition, we report a novel mislocalization pattern, with a bias towards the middle finger in the distal phalanges that reverses towards the thumb in the proximal phalanx, enriching the known distortions of the *superficial schema*. Overall, these findings provide novel insights into the functional organization of MBRs and their relationships with somatosensory information. Reducing the perceived body size through RSS could be useful in helping treat body image disturbance.

## Introduction

Mental body representations (MBRs) are critical for several, fundamental sensory abilities, such as localizing touches on our body surface and estimating body part’s size, as well as maintaining a coherent sense of bodily self. Indeed, tactile judgements are made by relating tactile inputs to MBRs, revealing a body-referencing of tactile perception (Serino & Haggard, 2010; Longo et al., 2010; de Haan & Dijkerman, 2020). The existence of different MBRs, related to specific sensorimotor functions, has been advocated by several models of MBRs. According to Serino & Haggard’s (2010) multilevel model of somatosensory perception and body representation, MBRs housed in posterior parietal areas are continuously updated by the activity of the primary somatosensory cortex (SI). Importantly, this model accounts for the well-established finding that MBRs do not reflect the morphology of body parts accurately, but - to some extent – their distorted SI representations (Tamè et al., 2021, Tamè & Longo, 2023). Longo & Haggard’s MBR model (Longo et al. 2010, 2015) also considers the involvement of specific MBRs in somatosensory processing, namely the *body model*, the *superficial schema* and the *body image*, and similarly proposes that they serve distinct purposes.

The *body model* is thought to be involved in perceiving metric properties of tactile stimuli like the tactile distance perceived between two points (Longo & Haggard, 2011), while the *superficial schema* is used for locating somatic stimuli on the body surface (Longo et al., 2015, Medina & Coslett, 2016). Whether inherent to the representations or a product of near-optimal Bayesian integration of somatosensory inputs (Peviani, Miller & Medendorp, 2024), perceptual distortions are consistently observed when assessing these MBRs. Indeed, Longo & Haggard (2010, 2011) observed that the hand is perceived wider and the fingers shorter when using indirect measures of hand size (e.g., the tactile distance perception and landmark localization tasks) indicating that the *body model* retains part of SI homuncular distortions. Similarly, using a tactile localization task on the hand’s dorsum, Mancini et al. (2011) reported distal biases towards fingertips and thumb, indicating that the *superficial schema* is also distorted. In contrast, the *body image* corresponds to a conscious and relatively accurate representation allowing for instance to estimate one’s hand’s shape and size accurately, as typically assessed through a template matching task (Longo & Haggard, 2010; 2011). All three MBRs were found to rely on the parietal cortex (Castellini et al., 2013; Klautke et al., 2023; Porro et al., 2007; Spitoni et al., 2010), though some findings suggest the *body image* additionally involves extra-parietal regions (Miyake et al., 2010; Castellini et al., 2013; Dary et al., 2023).

While it is widely accepted that somatosensory processes contribute to building and maintaining MBRs, their exact contribution to each remains unclear. Considering the different functions of MBRs, current models imply that different somatosensory processes may underlie different MBRs. As such, whether different MBRs are equally or differentially fed by tactile processing remains an open question. Answering this question has both theoretical and practical implications. Clarifying to which extent tactile inputs contribute to each MBR is crucial to refine current theoretical models and will allow a deeper understanding of MBRs’ interrelationship. In addition, this may open new avenues for the diagnosis and treatment of clinical conditions whereby MBRs are altered. To shed light on this issue, here we posit that if the three representations are similarly sustained by somatosensory activity, altering such activity would similarly affect them all. Alternatively, if they bear different relationships with somatosensory processes, some body representations could be affected differently.

One way to address this question consists in evaluating whether temporary modulating tactile information affects the different MBRs. Some pioneering studies investigated the effects of such modulation on a single MBR. For instance, seminal work by Gandevia & Phegan (1999) on the *body image* revealed that the body parts whose tactile inputs were temporarily reduced by anesthesia were perceived as bigger. More recently, Giurgola et al. (2019) showed that interfering with the activity of SI hand representation via repetitive Transcranial Magnetic Stimulation (rTMS) resulted in an overestimation of hand size. Thus, perception of body parts’ size via the *body image* seems to be modified when tactile inputs are severely reduced, or somatosensory processing altered. Moreover, TMS over the SI hand representation impairs tactile identification of stimulated fingers (Seyal, Siddiqui and Hundal, 1997), suggesting that interfering with SI function is also detrimental to the *superficial schema*.

While these studies indicate a tight link between somatosensory processes and MBRs, they typically investigated only one, or two MBRs. To gain deeper insights into the relationship between touch and MBRs, we assessed the impact of increasing tactile inputs on the three MBRs. To this aim, we leveraged the properties of repetitive somatosensory stimulation (RSS), known to temporarily improve tactile perception by modulating SI activity (Beste & Dinse, 2013). RSS consists in the passive stimulation of a given body part (typically the right index fingertip) for a prolonged period, to induce synchronized neuronal activations in the corresponding SI representation, resulting in its transient enlargement and in improved tactile perception (see Beste & Dinse, 2013 and Parianen Lesemann et al., 2015 for reviews). Here we used RSS as a tool to temporarily increase tactile inputs and investigate its effect on the three aforementioned MBRs, as assessed through three well-established paradigms: (i) the Template Matching Task (TMT) measuring the perceived size of the finger (involving the *body image*; Gandevia et al., 1999; Longo & Haggard, 2010; 2012a), (ii) the Tactile Distance Judgement Task (TDJT) assessing the distance between two tactile stimuli (involving the *body model* and the *superficial schema*; Longo & Haggard, 2011; Tamè et al., 2021), and (iii) the Tactile Localization Task (TLT) measuring the localization of tactile stimuli (involving the *superficial schema*; Mancini et al., 2011; Badde & Heed, 2016).

We found that RSS alters participants’ *body image* (TMT) without modifying the other MBRs/tasks, thus revealing that MBRs are not to be considered as equally dependent on somatosensory processes. Moreover, we found that increased tactile inputs (RSS) to the finger *reduces* the perceived finger size. This result, opposite to Gandevia and Phegan’s original report (1999) obtained following reduction of tactile inputs (anesthesia), suggests that the *body image* is sensitive to somatosensory modulation in both directions.

## Methods

### Participants

We included 33 healthy adults (27 women and 6 men; mean age ± SD: 22.8 ± 3.4 years). The sample size was determined by a power analysis using G*Power 3.1 (Faul et al., 2007) based on the available work on TMT (Ambron & Coslett, 2023) and TDJT (Taylor-Clarke et al., 2004; de Vignemont & Haggard, 2005; Tajadura-Jimenez et al., 2015), although the interventions in the TDJT studies were mainly visual and auditory. Their effect sizes were between medium (computed Cohen’s d= 0.5) and large (d= 0.8), their sample sizes ranging between 8 and 20. Our calculation showed that a sample size of 16 to 34 was required to detect a large to medium effect with 80% power using repeated measures ANOVAs. Participants were right-handed (Edinburgh handedness inventory (Oldfield, 1971), average score ± SD: 85.2 ± 15.6), without any neurological nor psychiatric disease, and with no history of injuries at the right index finger. They gave their written informed consent before participating and received compensation at the end of the study. Procedures were approved by the French ethics committee (CPP SUD EST IV n. ID RCB: 2010-A01180-39).

### Experimental timeline

A randomized, double-blind, sham-controlled design was used (Fig 1). All participants received the RSS and Sham interventions on the right index fingertip on two different days separated by a two-day washout period, since RSS effects on tactile perception are known to last up to 6 hours (Godde et al., 2000). Half of the participants received RSS first and the other half received Sham first, each participant being randomly assigned to either group. The effects of these interventions (RSS/Sham) on right index MBRs were investigated through Pre and Post testing sessions, including the TMT, the TDJT and the TLT delivered in a counterbalanced order. To verify RSS efficacy, tactile discrimination was also assessed before and after interventions through the 2PDT More details about experimental procedures are available in Supplementary materials.

**Fig 1.**
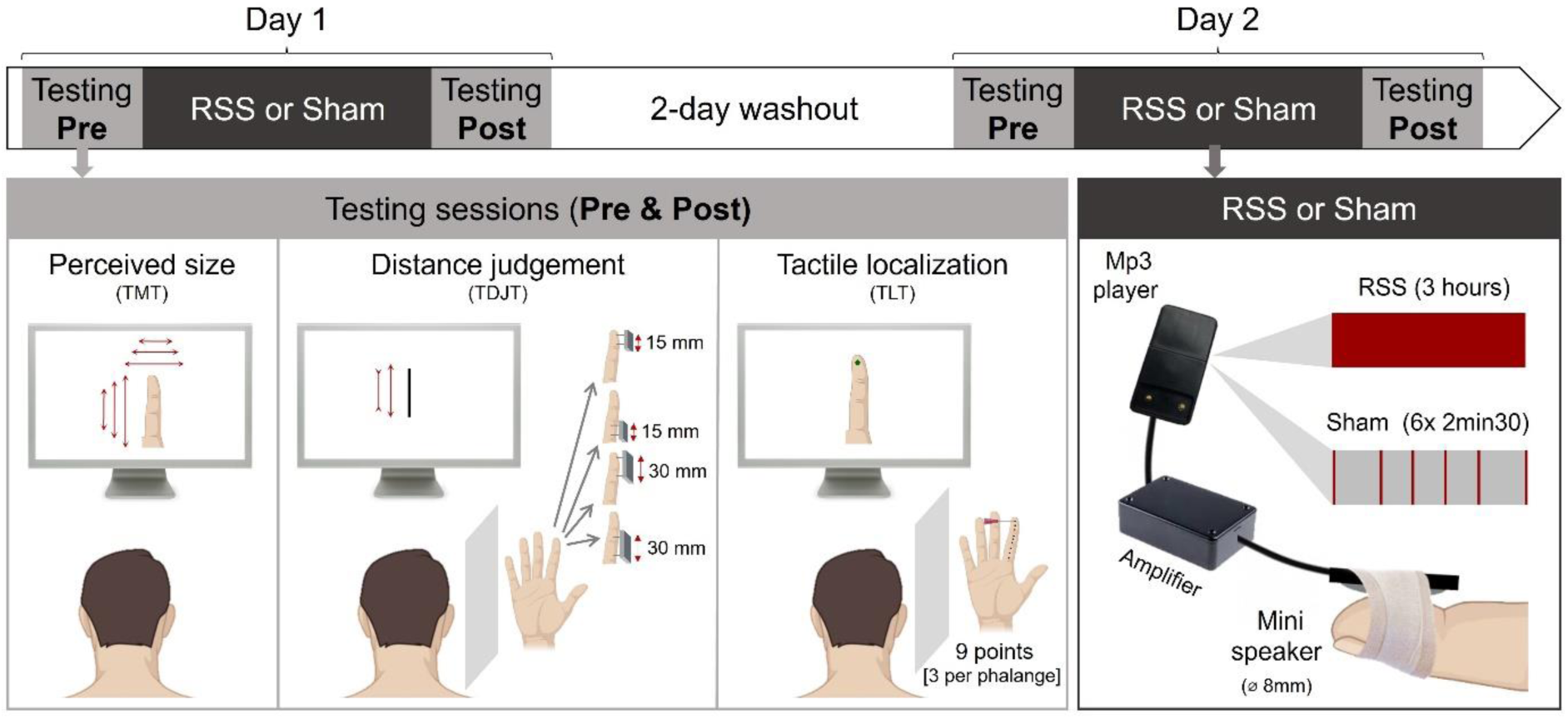
Experimental timeline and depiction of the tasks and interventions used. Participants received Sham and RSS on two different days (counter-balanced order). Before and after each intervention, perceived finger size, tactile distance judgement and tactile localization were assessed through the Template matching task (TMT), the Tactile Distance Judgement task (TDJT) and the Tactile localization task (TLT) respectively, in addition to the 2-point discrimination task (2PDT, not illustrated here).

Before the experiment, the participants’ right index finger was high-resolution color scanned, and the image was resized to match the real size of their finger using Photoshop ^®^, before being used in the TMT and TLT. The three tasks were implemented using MATLAB (MathWorks ^®^, version 2015b).

### Interventions (RSS & Sham)

The intervention protocols consisted in a 3-hour task-free mechanical stimulation on the right index finger. A small (8 mm diameter) mini loudspeaker (LSM-S20K, Ekulit) controlled by a mp3 player (Lenco Xemio-240 4GB) was taped to the right index fingertip (Fig 1). In the RSS protocol, this mini loudspeaker delivered brief (10ms) supra threshold tactile stimuli for 3 hours, with inter-stimulus intervals ranging from 100 to 3000ms and following a Poisson distribution (average stimulation frequency of 1Hz). The Sham protocol consisted in 15 minutes of tactile stimulation distributed across the 3 hours (i.e., 6 blocks of 2.5min each). Within each block, the mini loudspeaker delivered tactile stimuli with the same frequency and distribution as during RSS.

### Two-point Discrimination Task (2PDT)

Tactile acuity on the right index fingertip was assessed using the two-point discrimination task (2PDT) with a well-established procedure (Godde et al., 2000; Muret et al., 2014, 2016). Eight probes were presented on the volar surface of the fingertip: one with a single tip and seven with two tips separated by various distances (0.7, 1, 1.3, 1.6, 1.9, 2.2 and 2.5 mm). Each probe was tested 8 times in pseudo-randomized order, resulting in 64 trials per session. Tips were presented aligned to the longitudinal axis of the finger. Participants were blindfolded and asked to indicate whether they perceived “one” or “two” tips at each trial with the specific instruction of saying “two” only when the tips were clearly distinguishable. They did not receive feedback about their performance and had no time constraint to answer.

For each participant, the average of the verbal responses (“one” or “two”) was computed and the percentage of “two” responses was plotted as a function of the distance between the probes. The psychometric function was fitted with a binary logistic regression. From these fitted data, the PSE was determined for each session (Pre & Post) and intervention (Sham & RSS) (Fig 2A).

**Fig 2.**
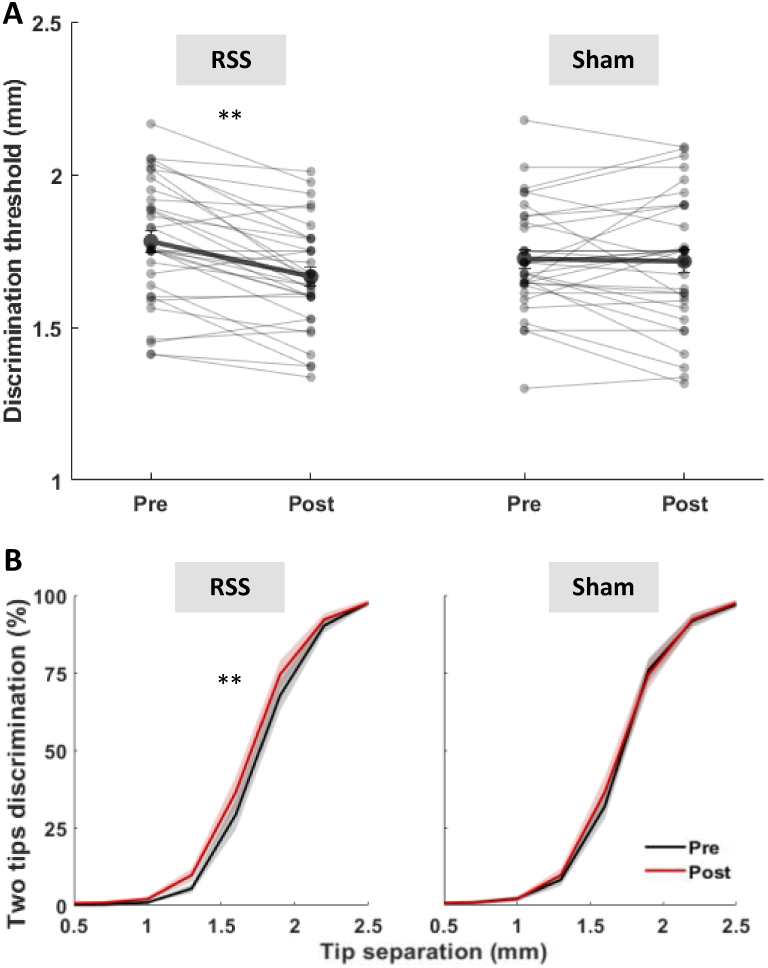
RSS successfully improved 2PDT threshold. (A) Individual (thin lines) and average (thick lines, ± SEM) discrimination thresholds obtained Pre and Post RSS or Sham interventions. (B) Mean psychometric curves of the 2PDT Pre (black) and Post (red) RSS or Sham interventions (mean ± SEM). ** p < 0.01 (paired t-tests, α_Bonf_ = 0.025).

### Template Matching Task (TMT)

Participants were seated in front of a computer screen (at a distance of 55 cm), tilted at 30° above the horizontal plane, with their left hand on a keyboard and their right hand hidden under the table, open with their palm facing upwards. At each trial, an image of their real index finger (real size, larger, or smaller) was displayed on the screen, and they were asked to judge whether this finger was smaller or larger than the actual size of their finger by pressing the corresponding keyboard buttons (+ or −). They were instructed to focus on their right index finger and to try to be as accurate as possible, without receiving feedback about their performance and without time constraints.

The stimuli presented to the participant were as follows: four enlarged, four reduced (with uniform area distortions of ±3%, ±6%, ±9%, ±12% relative to their actual finger size), and one real-sized image of their finger. Each stimulus was presented 12 times in randomized order for a total of 108 trials. For each participant, the percentage of responses corresponding to “image perceived larger” was plotted as a function of finger image distortion, and a psychometric function was fitted with a binary logistic regression (Statistica^TM^ Tibco ^®^, version 13.3). From these fitted data, the point of subjective equality (PSE), at which they perceived the image as big as their finger, was determined as the distortion threshold at which participants were at chance level (Fig 2A).

### Tactile Distance Judgement Task (TDJT)

Participants were seated in front of a computer screen (at a distance of 55 cm), with their unseen right hand lying supine on the table behind an occluding board. The experimenter touched the volar surface of their right index finger with 2 wooden rods simultaneously applied along the longitudinal axis, either within a single phalanx (i.e., rods spaced by 15 mm) or across two adjacent phalanges (i.e., rods spaced by 30 mm). For each distance, the rods were applied (for approximately 1 s) starting from 2 different locations: the base or the tip of the finger (Fig 1, middle lower panel). Each of the four conditions was repeated 10 times in a pseudo-randomized order, for a total of 40 trials. Participants were instructed to assess the tactile distance between the two rods by adjusting the length of a bar on the screen (pressing + and – buttons) to match the perceived distance between the two rods. They were instructed to focus on their right index finger sensation and to try to be as accurate as possible without any time constraint, and they did not receive feedback about their performance. The length of the bar as reproduced by participants was recorded and averaged for each session (Pre, Post), intervention (Sham, RSS), and position (base, tip).

### Tactile Localization Task (TLT)

Participants were seated in front of a computer screen, in the same configuration as in the TDJT. The experimenter touched the volar surface of their right index finger with a plastic von Frey monofilament of 5 g, at one of 9 different locations on the longitudinal midline of the finger, with 3 positions per phalanx: ¼, ½ and ¾ of the length of each phalanx. The locations were numbered from 1 to 9 with the 1^st^ location being the most proximal and the 9^th^ the most distal. Each location was touched 10 times in a pseudo-randomized order, for a total of 90 trials. The real-sized image of their own finger was displayed on a black screen in front of them, and they were asked to report on the image the exact location where they perceived the touch. To do so, participants moved with their left hand a green cursor on the screen to the desired location and validated their choice by pressing a button. They were instructed to focus on their right index finger sensation, and to try to be as accurate as possible without any time constraint. They did not receive feedback about their performance.

Both the judged (J) and real (R) locations were recorded as x and y coordinates of the picture displayed on the screen. The origin of the coordinate system was centered on each of the real locations, with the y-axis representing the longitudinal (proximo-distal) axis of the finger and the x-axis the bottom medio-lateral (ulnar-radial) axis of the finger. After normalizing the coordinates to each participant’s finger length, three measures were calculated for each of the 9 locations (see Supplementary Fig S1): (i) the Euclidean distance between the J and R locations was computed using the following formula: 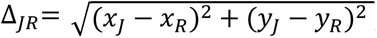, (ii) the polar angle between the JR vector and the x-axis at the R location was computed as: 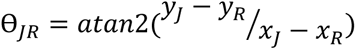, (iii) 95% confidence ellipses of the judged locations (with the mean judged location (x̅_J_, y̅_J_) being the center of the ellipse) were computed. The mean J-R distance and the mean J-R angle were used as a measure of constant error of tactile localization, while the mean area of the confidence ellipse was considered a measure of the variable error.

### Statistical analysis

Data were collected through Matlab (MathWorks ^®^, version 2015b). Data were missing for two participants in the TLT (for technical reasons), and two participants were removed from the analysis of the TMT given the impossibility of fitting the data with a binary logistic regression. In total, the number of participants included in each task are the following: n=33 in the 2PDT and TDJT, n=32 in the variable error of TLT and n=31 in the constant error of TLT and in the TMT, with 30 participants common to the 4 tasks. Outliers were defined as falling outside 3SD around the average. First, in tasks containing single trials (TDJT & TLT), outlier trials were identified (intra subjects). No outlier trials were found in either task. In the TLT, a few missed trials were removed, representing 0.03, 0.21, 0.35 and 0.14% of the data in Pre Sham, Post Sham, Pre RSS and Post RSS, respectively. After trials removal, a minimum of 7 trials out of 10 were left for each condition. Then, in all tasks, outliers were identified at the group level (inter subjects). When present, statistical analyses are reported in the results section both including and excluding these outliers, as this did not change the findings. Results in the text are expressed as mean ± SEM.

After verification of the normality (Shapiro-Wilk test), homoscedasticity (Levene’s test), and sphericity (Mauchly’s test) assumptions, repeated-measures ANOVAs (rmANOVAs) were conducted. When significant main effects or interactions were found, two-tailed t-tests were conducted with alpha levels Bonferroni-corrected for the number of tests performed (α_Bonf_). Effect sizes were computed using Cohen’s d (Cohen, 1988). When normality and/or homoscedasticity assumptions were not met, Friedman tests were conducted.

In the 2PDT and TMT, two-way rmANOVAs with the factors Intervention (Sham/RSS) and Session (Pre/Post) were used.

In the TDJT and TLT analyses, four-way rmANOVAs were used, with the common factors Intervention (Sham/RSS) and Session (Pre/Post); Position (Tip or Base of the finger) and Distance (15 mm or 30 mm) were also included for the TDJT analysis, and Position (n°1, n°2 or n°3 at each phalanx) and Phalanx (Proximal, Middle, Distal) for the TLT. Additionally, the difference between the real and the judged distances in the TDJT data was analyzed through two one-sample two-tailed paired t-tests (one for the 15 mm condition and the other for the 30 mm condition) and a two-tailed paired t-test to compare the two conditions. Finally, to determine the localization biases in the ulnar-radial (x) and proximo-distal (y) axes in the TLT data, the x and y components of the ΔJR vector were compared to zero (null bias) using one-sample two-tailed t-tests.

Except when specified otherwise, the threshold for statistical significance was set at p ≤ 0.05. Statistical analyses were performed through Jamovi (version 2.2.5). Complementary Bayesian t-tests (Pre vs. Post) are reported in Supplementary Table S1.

## Results

### Affecting tactile processes through RSS improves tactile perception

To assess whether RSS was successful in affecting somatosensory processing, we evaluated its impact on right D2’s 2PDT threshold. A two-way rmANOVA revealed a significant Intervention*Session interaction (*F*_(1,64)_ = 10.40, p = 0.002, *η²* = 0.02) arising from a significant decrease in perceptual thresholds after RSS (*t*_(32)_ = 5.14, p < 0.001, α_Bonf_ = 0.025, *d* = 0.89, 95% CI [0.49, 1.30]), while they remained stable after Sham (*t*_(32)_ = 0.19, p = 0.85, *d* = 0.03; Fig 2A). No significant difference was observed on the slope of the psychometric curves as checked through a Friedman test (*X²*_(3)_ = 0.34, p = 0.953; Fig 2B).

### RSS impacts the body image: the stimulated finger is perceived as smaller

We then assessed whether RSS had an impact on the finger size perception (i.e., *body image*) of the stimulated finger with the Template Matching Task (TMT). A two-way rmANOVA revealed a significant Intervention*Session interaction (*F*_(1,60)_ = 4.49, p = 0.038, *η²* = 0.01). Post-hoc paired t-tests showed that PSEs – expressed as the percentage of image distortion – were significantly smaller after RSS (*t*_(30)_ = 2.78, p = 0.009; α_Bonf_ = 0.025; *d* = 0.50, 95% CI [0.12, 0.87]; Fig 3A). On average, participants perceived their finger as −7.4 ± 4.1 % (mean ± SEM) smaller than it was before RSS. In contrast, no significant change was observed after the Sham intervention (*t*_(30)_ = −0.12, p = 0.909). As in the 2PDT, no significant difference in psychometric curves’ slopes was found across conditions (*X²*_F(3)_ = 2.27, p = 0.518; Fig 3B).

**Fig 3.**
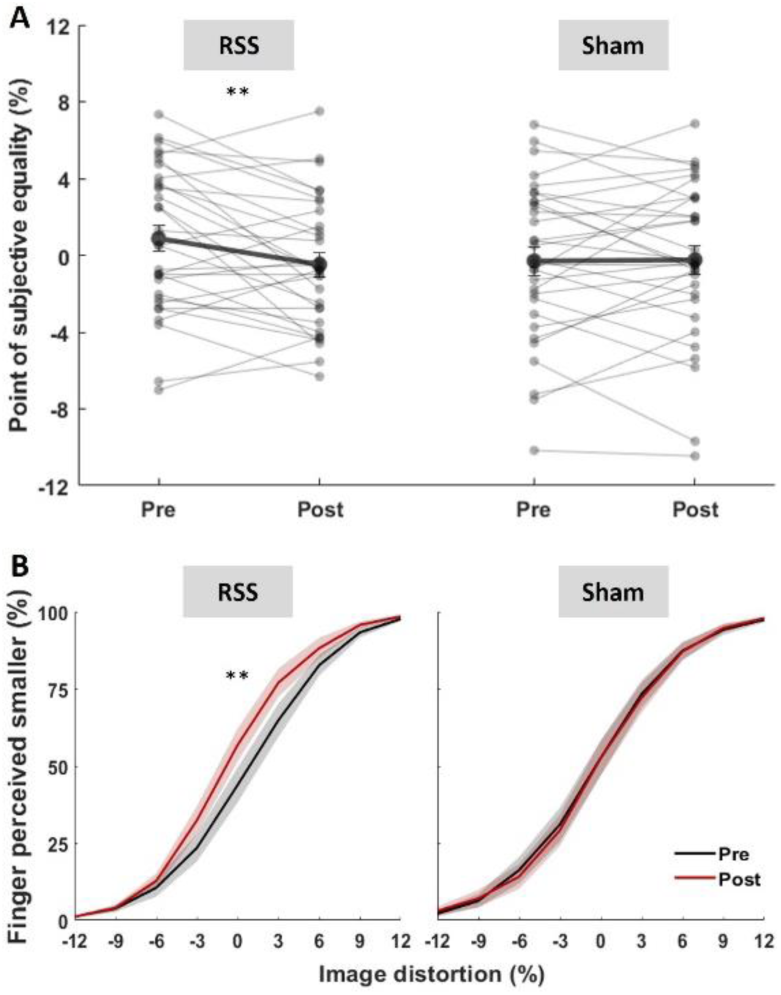
The stimulated finger is perceived as smaller after RSS but not after Sham. (A) Individual (thin lines) and average (thick lines, ± SEM) PSEs obtained Pre and Post RSS or Sham interventions. (B) Mean psychometric curves of the TMT Pre (black) and Post (red) RSS or Sham interventions (mean ± SEM). ** p < 0.01 (paired t-tests, α_Bonf_ = 0.025).

### RSS does not affect the body model and superficial schema of the stimulated finger

The Tactile Distance Judgment Task (TDJT) was used to assess any effect of RSS on the *body model* and *superficial schema*. A four-way rmANOVA showed no significant interaction between Interventions and Sessions (*F*_(1,64)_ = 3.35, p = 0.072, *η²* = 0.001), with no further interaction with the Distance and Position (both *F*_(1,64)_ ≤ 1.07, all p ≥ 0.35, *η²* = 0.001; see Fig 4A). Besides, distances were underestimated both in the within phalanx (15 mm) and the across phalanges (30 mm) conditions as revealed through two-tailed one sample t-tests comparing the perceived distances to the real distance (both *t* ≥ −11.30, both p values < 0.001, both *d* ≥ −1.96). No difference in underestimation between the two conditions was found through a two-tailed paired t-test (*t*_(32)_ = 0.96, p = 0.346). Finally, as expected from the higher density of mechanoreceptors (Johansson & Vallbo, 1979), distances (both 15 and 30 mm) were perceived significantly bigger (i.e., closer to the actual distance) at the tip than at the base of the finger (*F*_(1,64)_ = 53.16, p < 0.001, *η²* = 0.02). Similar results were obtained without outliers (n= 31; Supplementary Table S2).

**Fig 4.**
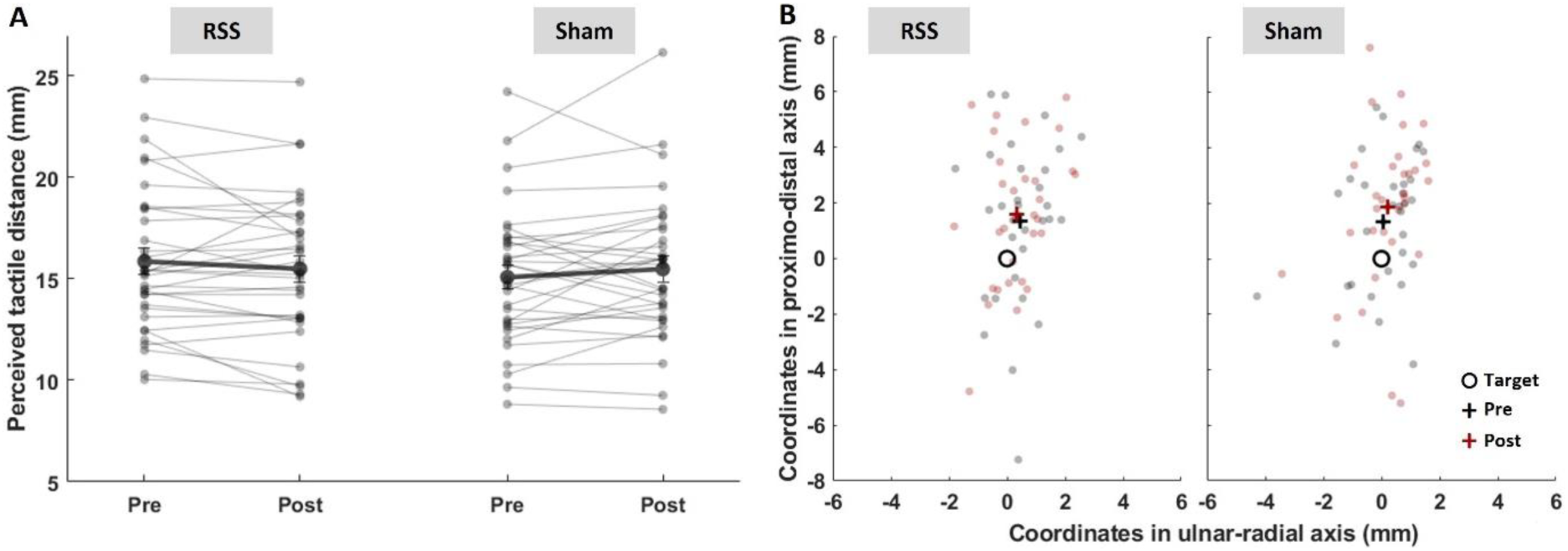
RSS does not alter the perceived tactile distance nor tactile localization. (A) Individual (thin lines) and average (thick lines, ± SEM) perceived tactile distances obtained Pre and Post RSS or Sham interventions. The data represented is averaged across the two positions (tip & base) and distances (15 & 30 mm). (B) Individual (dots) and average (crosses) judged positions relative to the target point (circle) Pre (black) and Post (red) RSS or Sham interventions. The data represented is averaged across the 9 points.

Another task used to assess the *superficial schema* is the Tactile Localization Task (TLT). Localization performance was compared to the actual target position for each of the 9 locations along the finger. Their x and y coordinates were used to compute two localization error estimates: the constant error and the variable error. While the constant error consists in the length (error magnitude) and orientation (angle relative to x axis; error direction) of a vector connecting the real and the averaged judged locations, the variable error is the dispersion of the judged locations measured as the area of the 95% confidence ellipse computed from the judged locations. Four-way rmANOVAs revealed no significant main effects nor interactions involving the factors Intervention (Sham/RSS) and Session (Pre/Post) for either the constant error measures (i.e., error magnitude and direction; all *F* ≤ 1.67, all p ≥ 0.199) nor the variable error measure quantified at each of the 9 locations (all *F* ≤ 2.21, all p ≥ 0.143; Fig 4B; detailed statistics in Supplementary Table S2). Similar results were obtained without outliers (n=26, 28 and 23 for error magnitude, direction and variable error respectively; Supplementary Table S2).

Overall, the results showed that RSS, which was effective in reducing the 2PDT threshold at the stimulated finger, significantly affected only one of the three measures we assessed, namely that considered to tap into the *body image*. No significant correlation was found between changes in TMT thresholds and changes in 2PDT thresholds (r = 0.16, p = 0.396).

### A systematic pattern of localization bias along the finger

Besides the effects of RSS, the rmANOVA ran on error magnitudes also revealed a significant difference between phalanges (*F*_(2,120)_ = 18.25, p < 0.001, *η²* = 0.10), arising from significantly shorter error magnitudes in the distal phalanx than in middle (*t*_(30)_ = 4.74, p < 0.001, α_Bonf_ = 0.017, *d* = 0.85) and proximal (*t*_(30)_ = 3.93, p < 0.001, α_Bonf_ = 0.017, *d* = 0.71) phalanges (Fig 5A). To further explore the pattern of these errors of localization per phalanx, we then compared the x and y coordinates of participants’ mean localization relative to the target (defined as the origin of the coordinate system). One-sample two-tailed t-tests comparing the x component (i.e., lateral error) to zero revealed a significant ulnar bias for the distal and middle phalanges (both *t*_(30)_ ≥ −4.18, both p < 0.001, α_Bonf_ = 0.017, both *d* ≤ −0.75; Fig 5B upper and middle panels), as well as a significant radial bias for the proximal phalanx (*t*_(30)_ = 3.34, p = 0.002, α_Bonf_ = 0.017, *d* = 0.60; Fig 5B lower panel). In the proximo-distal axis, one-sample two-tailed t-tests comparing the y component to zero revealed a significant proximal bias for the distal and middle phalanges (both *t*_(30)_ ≥ 4.68, both p < 0.001, α_Bonf_ = 0.017, both *d* ≥ 1.306; Fig 5B upper and middle panels). These results reveal a consistent bias in localization, with errors at the distal and middle phalanges directed towards the middle finger and the palm (proximo-ulnar bias), while errors at the proximal phalanx were directed towards the thumb (radial bias).

**Fig 5.**
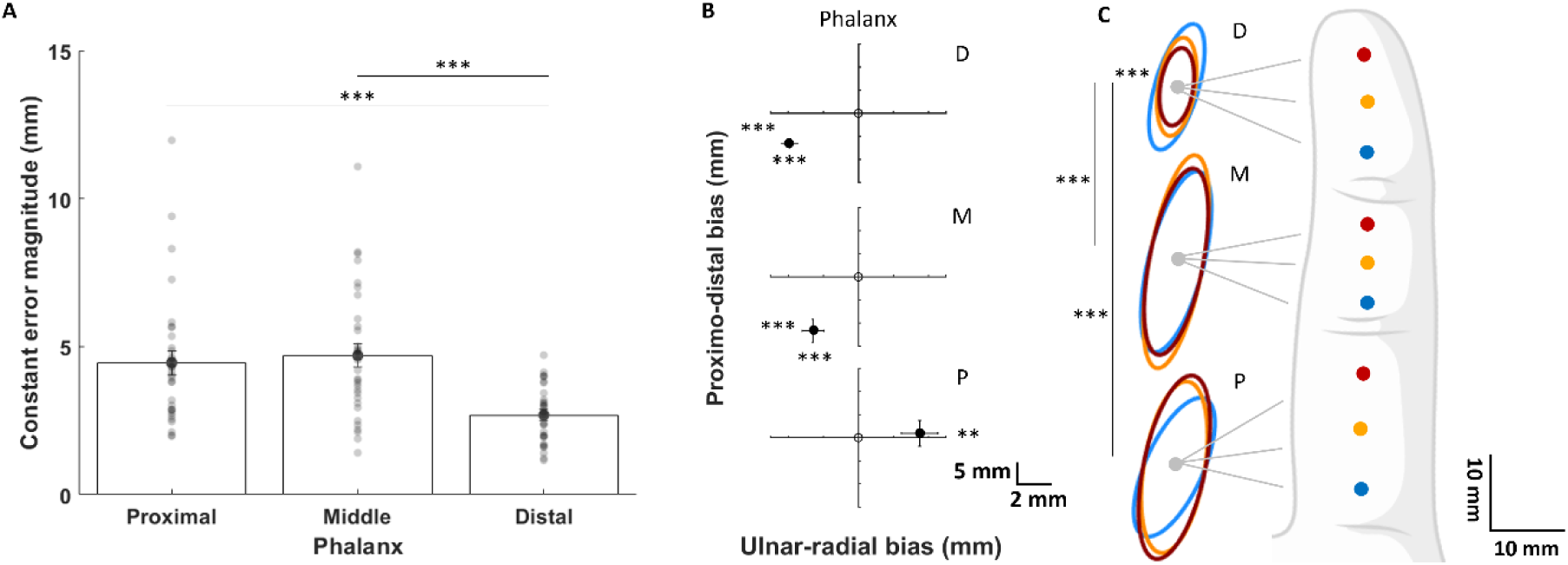
The TLT uncovered a pattern of perceptual bias along the finger. (A) Mean (± SEM) constant error magnitudes (length of error vectors) in the three phalanges. Errors were averaged across the 3 locations within each phalanx as they did not differ significantly(all *F*_(2,120)_ ≤ 1.19, all p ≥ 0.31). *** p < 0.001 (paired t-tests, α_Bonf_ = 0.017). (B) Mean (± SEM) difference between judged and target locations in ulnar-radial (x) and proximo-distal axis (y) for the proximal (P), middle (M) and distal (D) phalanges. *** p < 0.001; ** p < 0.01 (one-sample t-tests against zero, α_Bonf_ = 0.017). (C) Ellipses corresponding to the variable error for the blue, yellow and red target points within each of the three phalanges (proximal: P, middle: M, distal: D). Data is averaged across interventions (RSS and Sham) and sessions (Pre and Post) since those were not different (see main text). *** p < 0.001 (paired t-tests, α_Bonf_ = 0.0014).

Similarly to the constant error, the rmANOVA ran on the variable error revealed a significant main effect of Phalanx (*F*_(2,124)_ = 81.28, p < 0.001, *η²* = 0.15) arising from the areas of ellipses being significantly smaller at the distal than at the proximal (*t*_(31)_ = 8.39, p < 0.001, α_Bonf_ = 0.017, *d* = 1.48) or middle (*t*_(31)_ = 9.10, p < 0.001, α_Bonf_ = 0.017, *d* = 1.61) phalanges. A significant Phalanx*Point interaction was also observed (*F*_(4,248)_ = 8.31, p < 0.001, *η²* =0.015), with a notable additional gradient observed within the distal phalanx (Supplementary Table S3 for additional post-hoc comparisons, mostly replicating the main effect of Phalanges). Indeed, the most distal ellipse (red dot on phalanx D in Fig 5C) was significantly smaller than the middle (yellow dot on phalanx D: *t*_(31)_ = 3.94, p < 0.001, α_Bonf_ = 0.0014, *d* = 0.70), itself significantly smaller than the proximal one (blue dot on phalanx D: *t*_(31)_ = 4.45, p < 0.001, α_Bonf_ = 0.0014, *d* = 0.79). Similar results were obtained without outliers (n=26 and 23 for error magnitude and variable error respectively; Supplementary Table S2).

## Discussion

The present study aimed to elucidate the link between MBRs and their supposedly shared somatosensory basis. To this aim, we investigated the effect of increasing tactile inputs via RSS – known to reduce the 2PDT thresholds of the stimulated finger by modulating SI activity (Pleger et al., 2001, 2003) – on three MBRs of the same finger. Following either RSS or Sham intervention on the index finger, we assessed the *body image*, as probed through a Template Matching Task, the *body model*, as probed through a Tactile Distance Judgement Task, and the *superficial schema*, as probed through the Tactile Distance Judgement Task and a Tactile Localization Task. We first ascertained RSS efficacy by replicating the expected finding of improved 2PDT performance at the index finger (Godde et al., 2000; Pleger et al., 2001, 2003, Muret et al., 2014, 2016). We then reported a reduction of perceived finger size following RSS and no change in tactile distance judgement or tactile localization. These results suggest that increasing somatosensory inputs alters the *body image.* Instead, RSS did not alter the *body model* or the *superficial schema* in either direction.

### Increasing tactile inputs alters the body image

The *body image* is known to be quite accurate (Longo & Haggard, 2010; Longo et al., 2010, 2015). Our findings corroborate this notion, as we found a baseline distortion of only 0.3% (0% = real size). After RSS, whose efficacy was confirmed by the tactile improvement in the 2PDT, participants perceived their finger as being smaller than before. This suggests that increasing inputs through RSS altered the *body image*. This finding is consistent with the results of Gandevia & Phegan (1999) and Ambron & Coslett (2023) where the opposite modulation of inputs (i.e., reduction of inputs through anesthesia) increased the perceived size of the anesthetized body part. Altering tactile inputs thus seems to directly and bidirectionally impact the *body image*. One possible mechanism underlying the reduced body image of the finger may relate to the well-established effects of RSS on SI. By co-activating several skin receptive fields on the fingertip repeatedly for a protracted period of time, RSS has been shown to induce an enlargement of the stimulated finger’s representation in SI – increasing the neuronal resources available to process inputs – through long-term potentiation-like plasticity (Godde et al., 1996; 2000; Pleger et al., 2001, 2003). The *RSS-induced reduction* of *body image* through an increase of SI finger area activity is coherent with the *anesthesia-induced increase* of *body image* (Gandevia & Phegan, 1999; Ambron & Coslett, 2023) possibly through an attenuation of SI finger area activity. Indeed, it has been reported that anesthetizing the index finger through pharmacological nerve block attenuates the activity of the SI finger area, as observed in fMRI (Wesselink et al., 2022).

### Increasing tactile inputs does not modify the *body model* and the *superficial schema*

Unlike the *body image*, the *body model* and *superficial schema* are known to be distorted. Previous work reported an underestimation of the length of the fingers, together with a widening of the hand’s width (Longo & Haggard, 2010; Mancini et al., 2011; Coelho et al., 2017). Our results are in agreement with these distortions as we found an underestimation of tactile distances, as well as a proximal bias in tactile localization for both the distal and middle phalanges and no proximo-distal bias in the proximal phalanx. Besides, our findings are in keeping with well-established characteristics related to variations in mechanoreceptors density along the finger. Indeed, mechanoreceptors density follow a gradient along the finger (Johansson & Vallbo, 1979; Ciano & Beatty, 2022) with the higher density at the fingertip being associated with higher spatial discrimination (Johansson & Vallbo, 1979), resulting in distances being perceived as bigger (Weber, 1996) and tactile localization being more accurate (Yoshioka et al., 2013). Besides replicating the distances perceived bigger at the fingertip, localization performance showed also smaller constant and variable errors in the distal phalanx (with an additional gradient of variability within the distal phalanx) indicating higher localization accuracy and precision.

When it comes to RSS effects, in contrast to the *body image*, RSS did not alter either the *body model* or the *superficial schema*. One should consider that the tasks underlying these MBRs might be less sensitive to the increase in tactile inputs induced by RSS. The fact that the *body image* and the *body model*/*superficial schema* may be differentially affected is consistent with the results of Mergen et al’s study (2018), whereby patients suffering from anorexia nervosa – known to overestimate their body size, especially on the abdomen region (Keizer et al., 2012) – displayed preserved tactile localization ability on the abdomen. This converges with our findings in indicating that the *body image* can be affected selectively with respect to the *superficial schema*.

Other sensory manipulations have been previously reported to alter the *superficial schema*. Taylor-Clarke et al. (2004) found that visually magnifying the forearm and minifying the hand (i.e., changing their visual size) for 1h, decreased the well-established bias resulting in a bigger perceived distance on the finger than on the forearm. In other words, when the hand looks smaller, tactile distance is also perceived smaller as compared to when the hand appearance is veridical. Similarly, increased perceived tactile distance was observed when illusorily elongating the finger or the arm using either tendon stimulation (de Vignemont & Haggard (2005)) or a multisensory audio-tactile task (Tajadura-Jiménez et al. (2015)). Overall, these findings concur in showing that altering the perceived size of a body part (but not necessarily its body image) can affect the perceived distance between two points applied on this body part. However, it is worth noting that these manipulations consisted essentially in visual, proprioceptive and auditory illusions, making them less comparable to alterations of tactile inputs as implemented in the present work or following anesthesia. Together with our results, and knowing that MBRs are built based on both somatosensory (tactile and proprioceptive) and visual information (Bremner, 2016; de Klerk et al., 2021), we suggest that the *body model* and *superficial schema* may be less vulnerable to tactile manipulations than the *body image*. This may reflect either their relative ‘immunity’ to changes in tactile inputs, or their higher susceptibility to sensory correction: e.g., the altered tactile information could be compensated for by the intact proprioceptive and visual information.

Indeed, the *body model* and the *superficial schema* may generally be more rigid (i.e., their distortions seem less susceptible to change) than the *body image*. As shown by Longo & Haggard (2010, 2011) and Mancini et al. (2011), the distortions of the *body model* and *superficial schema* – as assessed through tactile size perception, landmark localization, and tactile localization tasks – are still found in different finger postures and hand orientations. Additionally, recent works (Bassolino & Becchio, 2023; Longo, 2023; Coelho & Gonzalez, 2024) seem to indicate that localizing tactile stimuli on the skin could require a correction factor. In this respect, the RSS-induced change in SI information received by the *superficial schema* and *body model* might have been corrected by this factor. This factor might apply only to body representations that are distorted and more action control-related (Bassolino & Becchio, 2023; Longo, 2023; Coelho & Gonzalez, 2024) and not (or less so) to body representations that are accurate and more perception-related such as the *body image* (Dijkerman & de Haan, 2007).

### Changes in *body image* are not linked to changes in perceptual thresholds

Regarding the potential relationship between the *body image* and 2PDT threshold changes, the lack of correlation suggests that these changes are not linearly related. The concomitant reduction of 2PDT threshold and *body image* size is coherent with the higher 2PDT thresholds found in patients exhibiting a bigger body image size (i.e., anorexia nervosa and chronic regional pain syndrome patients; Gadsby, 2017; Moseley et al., 2005, Keizer et al., 2012; Pleger et al., 2006), as compared to healthy populations. Yet, some studies reported an inverse relationship between such changes, with (i) a reduction of 2PDT thresholds and an increase of body image size following anesthesia (Ambron & Coslett, 2023), or (ii) an increase of 2PDT thresholds and a reduction of body image size following tendon vibration illusion (D’amour et al., 2015). Thus, the direction of change of 2PDT threshold does not seem to depend on the direction of change of the *body image* size.

### A novel pattern of localization bias along the finger

Besides replicating known distortions in MBRs and providing evidence for their differential sensitivity to modulation and reliance on tactile inputs, our results also bring new insights regarding MBRs. Intriguingly, we observed a specific pattern of ulnar-radial localization bias across phalanges, with the localization biased towards the thumb in the proximal phalanx and towards the middle finger in the more distal phalanges. Although localization within a single finger has rarely been investigated (Miller et al., 2022), studies assessing localization at the whole hand scale, targeting the tips and bases of fingers on the palmar surface, did not report such an ulnar-radial bias (Longo et al., 2012b, Dupin et al., 2022). This discrepancy may arise from the fact that they targeted the fingertip and the skin crease at the base of the finger, that can be considered as “landmarks”, while we targeted points away from creases/joints, equally distributed along the finger. It could also be due to postural differences as in our study, the hand was in a “natural” posture with fingers not splayed (abducted) nor pressed together (adducted), while in previous studies fingers were maintained abducted. A postural effect on tactile localization has been observed with localization on splayed fingers resulting in a wider hand representation than adducted fingers (Longo, 2015). Nevertheless, the radial bias of the proximal phalanx we newly report here seems in keeping with the radial bias found on the palm (Culver, 1970). Altogether, our findings may reflect a bias towards adjacent fingers, at locations where informative tactile (co-)stimulation across fingertips is more likely to occur due to postural and movement synergies. Indeed, in a natural hand posture, the proximal phalanx of the index finger contacts both the middle finger and the thumb while the other two phalanges contact only the middle finger. This finding raises new questions about the role of tactile “synergies” in the distortions observed at the level of the *superficial schema*.

### Updating the theoretical framework of MBRs

These findings help revising the model of MBRs, in particular with respect to their relationships with the tactile inputs (Fig 6). Indeed, the relationship between tactile inputs and the *body image* appears different from those linking them to the *superficial schema* and *body model*. While RSS is known to affect SI – nourishing all MBRs housed in the parietal cortex – this effect alone cannot account for the whole pattern of results observed in this study. The way SI exerts its modulatory effects differently on MBRs may either depend upon different sub-regions underpinning the *body imag*e and the other two MBRs, and/or upon the MBRs being linked to SI in a different way. Alternatively, beyond the parietal cortex, RSS-induced effects might spread to other areas exclusively involved in the *body image*. Indeed, the *body image* has been shown to involve also occipital, temporal and frontal areas (Miyake et al., 2010; Castellini et al., 2013; Dary et al., 2023) that have not been identified in the other two MBRs, predominantly relying on parietal cortex (Klautke et al., 2023; Porro et al., 2007; Spitoni et al., 2010). While future work will help disentangling these alternatives, our study provides novel evidence on the neglected link between MBRs and somatosensory processes, and paves the way to novel clinical applications for treatment of pathological MBR conditions.

**Fig 6.**
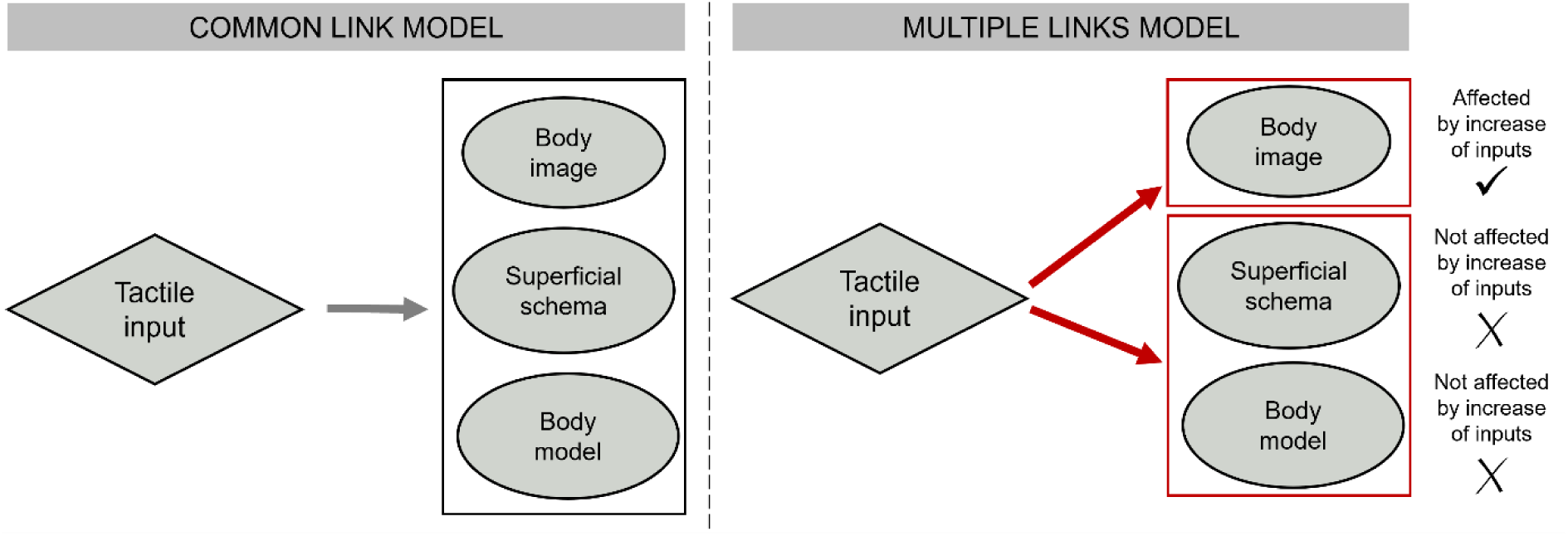
Update of the model of MBRs-tactile input relationship. With regard to the relationship between tactile inputs and MBRs, there was no known distinction between the three MBRs. Following our results showing that RSS affects selectively the *body image*, we propose a model revision that splits the tactile input – MBRs relationship into (at least) two distinct relationships: (i) tactile input – *body image* and (ii) tactile input – *superficial schema*/*body model*.

## Conclusion

We provide evidence that converge with previous work in indicating that the *body image* is bi-directionally susceptible to changes following a temporary modulation of tactile inputs. Our findings also indicate that MBRs, even if all nourished by tactile afferents through SI, are not affected in the same way by increasing tactile information. We suggest that the *body model* and *superficial schema* may be more rigid and less affected by modulation of tactile inputs. Importantly, this study provides a proof-of-concept that a simple non-invasive and effortless tactile stimulation can alter the *body image* in the direction of a reduction of the perceived body size, which could translate into rehabilitative strategies to help treat *body image* disturbance, frequently occurring in eating disorders (anorexia nervosa, bulimia).

## Acknowledgments

This work was supported by the Agence Nationale de la Recherche (ANR-19-CE37-0005) BLIND_TOUCH to AF and LEM, the Paul Bennetot Foundation (DL/VH 2018206) to AF, the Fondation pour la Recherche Médicale Doctoral Fellowship ECO202006011658 to MA-S, the Neurodis Foundation to AF and MA-S & the Federation pour la Recherche sur le Cerveau to AF and DM. We thank Frederic Volland for his help constructing the experimental setup, Valeria Ravenda and Isabella Bertoncini for helping with data collection, and Sonia Alouche, Sandra Chinel, Florence Leger and Celia Farge for administrative support.

## Authors contributions

MA-S, SM, LEM, MRL, DM and AF conceived the study and designed the experiment. L.E.M wrote some scripts for the experimental tasks. EK and RS designed the Sham stimulation protocol. MA-S and SM collected and analyzed the data. MA-S, DM and AF wrote the manuscript with inputs from all co-authors. All authors approved of the final submission.

