## Supplemental Material for "Repetitive Somatosensory Stimulation Shrinks The Body Image"

### Experimental procedures:

To ensure the same skin locations were tested in Pre and Post sessions for the TDJT and TLT, while preventing participants from seeing the marks on their finger, the locations were drawn on the skin with invisible ink which was made visible when lighted with UV light (during the experimental sessions only, hidden from the participant's sight). In order to make the ink clearly visible to the experimenter, both tasks were conducted in a darkened room. The TMT was also conducted in darkness to have a similar experimental environment.

Instructions during RSS & Sham stimulations:

During both RSS and Sham stimulations, participants were instructed not to attend to the stimulation and to continue with their daily activities, but to avoid intensive use of their fingers (e.g., typing on a keyboard) to avoid major concomitant sensorimotor activity.

### Details about the tasks:

2PDT:

Each probe was tested 8 times in pseudo-randomized order, resulting in 64 trials per session. To prevent desensitization of the skin area under assessment due to repeated indentation, short breaks were allowed every 20-30 trials. After allowing the participant to feel the extreme distances (i.e., 0 and 2.5 mm) a familiarization phase was performed (in which each probe was tested 4 times in pseudo-randomized order, for a total of 32 trials). Then, in order to try to achieve a stable baseline performance, two sessions (S1 & S2) separated by a 20-minute break were conducted, as performed in previous studies (Muret et al., 2014; 2016).

Thresholds obtained at S1 and S2 were statistically analyzed for stability with a rmANOVA with the factors Intervention (Sham/RSS) and Time (S1/S2). No main Time effect ($F(1,62) = 0.26$, $p = 0.609$, $\eta^2 = 0.001$) nor interaction ($F(1,62) = 0.02$, $p = 0.883$, $\eta^2 = 0.000$) were found, indicating that discrimination thresholds were stable at baseline. The data of S2 were considered as the Pre session data.

TMT:

Each stimulus was presented 12 times in randomized order for a total of 108 trials. A familiarization session consisting in 9 trials (each stimulus presented once) was conducted before starting the experiment.

TDJT:

The location at the base of the finger was placed at a distance of ¼th of the length of the proximal phalanx from the crease separating the finger from the palm, while the location at the tip of the finger was placed at a distance of ¼th of the length of the distal phalanx from the tip of the finger. Each of the four possible conditions (i.e., 15mm-base, 15mm-tip, 30mm-base, 30mm-tip) was repeated 10 times in a pseudo-randomized order, for a total of 40 trials.

On the screen, the initial size of the bar was either of 1 mm or 45 mm in a pseudo-randomized order.

TLT:

Each of the 9 locations along the finger was touched 10 times in a pseudo-randomized order, for a total of 90 trials. From these, the constant and variable localization errors were computed at each point as described in Fig S1.

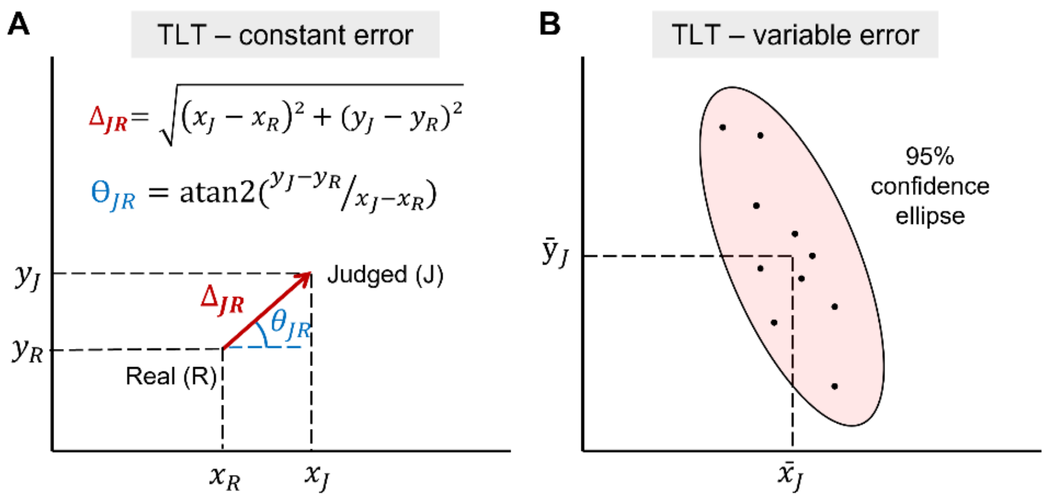


**Fig S1. Representation of the outcome measures of interest computed at the individual level in the TLT task.** (A) To determine the constant error, the difference between the judged (J) and real (R) locations as well as the angle between the JR vector and the x-axis at the R location were computed from their x and y coordinates. (B) To determine the variable error, the 95% confidence ellipses were computed from the judged locations (black dots), and their area was extracted (colored in pink here).

### Supplementary Table S1. Results of Bayesian paired sample t-tests ($BF_{incl}$)

| Task | Sham Pre vs Sham Post | RSS Pre vs RSS Post |
|---|---|---|
| 2PDT | 0.189 | 1608.864 |
| TMT | 0.193 | 4.759 |
| TDJT | 0.386 | 0.419 |
| TLT – constant error magnitude | 0.418 | 0.213 |
| TLT – constant error direction | 1.773 | 0.432 |
| TLT – variable error | 1.247 | 0.198 |

### Supplementary Table S2. Results of statistical analyses with and without the inter subjects outliers

| | | | | All subjects included | | | | Without outliers | | | |
|---|---|---|---|---|---|---|---|---|---|---|---|
| Task | Variable | Test | Effect | Statistic (F or t) | df | p | η² or d | Statistic (F or t) | df | p | η² or d |
| TDJT | Judged distance (n= 33) | rmANOVA | Group*Session | 3.35 | 1; 64 | 0.072 | 0.001 | 3.151 | 1; 60 | 0.081 | 0.001 |
| | | | Group*Session*Position | 1.065 | 1; 64 | 0.306 | 0.000 | 1.891 | 1; 60 | 0.174 | 0.000 |
| | | | Group*Session*Distance | 0.902 | 1; 64 | 0.346 | 0.000 | 1.072 | 1; 60 | 0.305 | 0.000 |
| | | | Position | 53.16 | 1; 64 | <0.001 | 0.02 | 72.137 | 1; 60 | <0.001 | 0.02 |
| | | One-sample t-test (judged vs real) | Judged vs real 15mm cond | -10.7 | 32 | <0.001 | -1.86 | -14.2 | 30 | <0.001 | -2.55 |
| | | | Judged vs real 30mm cond | -11.3 | 32 | <0.001 | -1.96 | -13.7 | 30 | <0.001 | -2.46 |
| | Real-Judged difference (n= 33) | Paired t-tests | 15 vs 30 mm | 0.96 | 32 | 0.346 | 0.167 | 1.51 | 31 | 0.141 | 0.267 |
| TLT | Constant error magnitude (n= 31) | rmANOVA | Group*Session | 1.671 | 1; 60 | 0.201 | 0.001 | 0.262 | 1; 50 | 0.611 | 0.000 |
| | | | Group*Session*Phalanx | 1.238 | 2; 120 | 0.294 | 0.001 | 1.336 | 2; 100 | 0.268 | 0.001 |
| | | | Group*Session*Point | 0.018 | 2; 120 | 0.982 | 0.000 | 0.030 | 2; 100 | 0.971 | 0.000 |
| | | | Phalanx | 18.251 | 2; 120 | <0.001 | 0.102 | 15.799 | 2; 100 | <0.001 | 0.097 |
| | | Post-hoc t-test (Phalanx) ** | Distal vs Middle | 4.735 | 30 | <0.001 | 0.850 | 3.782 | 25 | <0.001 | 0.742 |
| | | | Distal vs Proximal | 3.934 | 30 | <0.001 | 0.707 | 4.201 | 25 | <0.001 | 0.824 |
| | Constant error direction (n= 31) | rmANOVA | Group*Session | 0.443 | 1; 60 | 0.508 | 0.000 | 1.328 | 1; 54 | 0.254 | 0.001 |
| | | | Group*Session*Phalanx | 1.638 | 2; 120 | 0.199 | 0.001 | 2.300 | 2; 108 | 0.105 | 0.002 |
| | | | Group*Session*Point | 1.533 | 2; 120 | 0.220 | 0.001 | 0.876 | 2; 108 | 0.419 | 0.001 |
| | Variable error (n= 32) | rmANOVA | Group*Session | 2.205 | 1; 62 | 0.143 | 0.001 | 0.306 | 1; 44 | 0.583 | 0.000 |
| | | | Group*Session*Phalanx | 0.218 | 2; 124 | 0.804 | 0.000 | 1.989 | 2; 88 | 0.143 | 0.002 |
| | | | Group*Session*Point | 0.369 | 2; 124 | 0.693 | 0.000 | 0.936 | 2; 88 | 0.396 | 0.001 |
| | | | Phalanx | 81.281 | 2; 124 | <0.001 | 0.152 | 64.656 | 2; 88 | <0.001 | 0.183 |
| | | | Phalanx*Point | 8.310 | 4; 248 | <0.001 | 0.015 | 9.658 | 4; 176 | <0.001 | 0.030 |
| | | Post-hoc t-test (Phalanx) ** | Distal vs Middle | 9.102 | 31 | <0.001 | 1.609 | 9.646 | 22 | <0.001 | 2.011 |
| | | | Distal vs Proximal | 8.385 | 31 | <0.001 | 1.482 | 9.156 | 22 | <0.001 | 1.909 |
| | | Post-hoc t-test (Point*Phalanx) *** | Within distal : D vs M | 3.936 | 31 | <0.001 | 0.696 | 3.079 | 22 | 0.005 | 0.642 |
| | | | Within distal : M vs P | 4.448 | 31 | <0.001 | 0.786 | 3.087 | 22 | 0.005 | 0.644 |

* Without outliers: Judged distance (n=31); Real-Judged difference (n=32); Constant error magnitude (n=26); Constant error direction (n=28); Variable error (n=23)

** $\alpha_{Bonf} = 0.017$ *** $\alpha_{Bonf} = 0.006$

**Supplementary Table S3. Post-hoc tests following the significant Phalanx*Point interaction in the variable localization error with and without the inter subjects outliers**

| | | | All subjects included (n=32) | | | | Without outliers (n=23) | | | |
|---|---|---|---|---|---|---|---|---|---|---|
| Phalanx | Point 1 | Point 2 | Statistic (t) | df | p* | d | Statistic (t) | df | p* | d |
| Proximal | Proximal | Middle | -1.662 | 31 | 0.107 | -0.294 | -2.807 | 22 | 0.010 | -0.585 |
| | Proximal | Distal | -1.753 | 31 | 0.089 | -0.310 | -3.005 | 22 | 0.007 | -0.627 |
| | Middle | Distal | -0.997 | 31 | 0.326 | -0.176 | -0.993 | 22 | 0.331 | -0.207 |
| Middle | Proximal | Middle | -1.094 | 31 | 0.282 | -0.193 | -2.041 | 22 | 0.053 | -0.425 |
| | Proximal | Distal | -0.209 | 31 | 0.836 | -0.037 | -1.131 | 22 | 0.270 | -0.236 |
| | Middle | Distal | 1.023 | 31 | 0.314 | 0.181 | 0.656 | 22 | 0.519 | 0.137 |
| Distal | Proximal | Middle | 4.448 | 31 | <0.001 | 0.786 | 3.087 | 22 | 0.005 | 0.644 |
| | Proximal | Distal | 5.287 | 31 | <0.001 | 0.935 | 3.897 | 22 | <0.001 | 0.813 |
| | Middle | Distal | 3.936 | 31 | <0.001 | 0.696 | 3.079 | 22 | 0.005 | 0.642 |

* $\alpha_{Bonf} = 0.006$
